# Nuclear TARBP2 drives oncogenic dysregulation of RNA splicing and decay

**DOI:** 10.1101/389213

**Authors:** Lisa Fish, Hoang C.B. Nguyen, Steven Zhang, Myles Hochman, Brian D. Dill, Henrik Molina, Hamed S. Najafabadi, Claudio Alarcon, Hani Goodarzi

## Abstract

Post-transcriptional regulation of RNA stability is a key step in gene expression control. We describe a regulatory program, mediated by the double-stranded RNA binding protein TARBP2, that controls RNA stability in the nucleus. TARBP2 binding to pre-mRNAs results in increased intron retention, subsequently leading to targeted degradation of TARBP2-bound transcripts. This is mediated by TARBP2 recruitment of the m^6^A RNA methylation machinery to its target transcripts, where deposition of m^6^A marks influences the recruitment of splicing regulators, inhibiting efficient splicing. Interactions between TARBP2 and the nucleoprotein TPR then promote degradation of these TARBP2-bound transcripts by the nuclear exosome. Additionally, analysis of clinical gene expression datasets revealed a functional role for this TARBP2 pathway in lung cancer. Using xenograft mouse models, we find that TARBP2 impacts tumor growth in the lung, and that this function is dependent on TARBP2-mediated destabilization of ABCA3 and FOXN3. Finally, we establish the transcription factor ZNF143 as an upstream regulator of TARBP2 expression.

**RESEARCH HIGHLIGHTS:**

- The RNA-binding protein TARBP2 controls the stability of its target transcripts in the nucleus
- Nuclear TARBP2 recruits the methyltransferase complex to deposit m^6^A marks on its target transcripts
- TARBP2 and m^6^A-mediated interactions with splicing and nuclear RNA surveillance complexes result in target transcript intron retention and decay.
- Increased TARBP2 expression is associated with lung cancer and promotes lung cancer growth *in vivo*.
- The transcription factor ZNF143 drives oncogenic TARBP2 upregulation in lung cancer.

## INTRODUCTION

Post-transcriptional regulation of gene expression plays a major role in normal cell physiology and human diseases. The major molecular processes that are responsible for RNA turnover in the cytoplasm and the nucleus have been described in detail (Kilchert et al., 2016; Nasif et al., 2017). However, the regulatory programs that feed into these pathways to modulate transcript stability, and their collective role in shaping the cellular gene expression landscape, remain largely unexplored. Recently, targeted intron retention has been described as a mechanism that modulates RNA degradation (Wong et al., 2016). In this pathway, transcripts with retained introns that are exported to the cytoplasm may be degraded by nonsense-mediated decay factors, or may be targeted by the nuclear RNA surveillance machinery prior to export. This latter mechanism has been reported for individual transcripts (Bergeron et al., 2015), as well as for controlling gene expression patterns during neuron development (Yap et al., 2012). However, the upstream regulatory programs that are involved in these nuclear RNA decay processes remain largely unknown. Here, we report the discovery and characterization of one such post-transcriptional regulatory network that functions in the nucleus to govern RNA stability.

Recently, we described a previously unknown regulatory pathway in which the double-stranded RNA binding protein TARBP2 binds and destabilizes its target transcripts through an unknown mechanism (Goodarzi et al., 2014). In this study, we demonstrate that TARBP2 functions in the nucleus and modulates the stability of its regulon by influencing the rate of intron retention in its targets. Our findings reveal that nuclear TARBP2 recruits the RNA methylation machinery, resulting in local m6A-mediated remodeling of splicing factors and impeding efficient processing of its target transcripts. RNA molecules with retained introns are then dispatched for degradation through interactions between TARBP2 and the nuclear RNA surveillance complex. This novel regulatory program uncovers an interaction network between RNA modification, processing, and surveillance machineries, and reveals how they can function in concert to modulate the expression of a large regulon.

Our findings also highlight the emergence of RNA methylation as a major factor in post-transcriptional regulation of gene expression in the nucleus. The prevalent internal RNA modification mark N(6)-methyladenosine (m^6^A) has been reported to play a role in regulating most facets of the RNA life cycle, including regulation of pre-mRNA splicing, mRNA stability, and mRNA translation (Lin et al., 2016; Liu et al., 2015; Wang et al., 2015, 2014; Xiao et al., 2016; Zhao et al., 2014). Recently, work by us (Alarcón et al., 2015) and others (Ke et al., 2017; Knuckles et al., 2017) has established that m^6^A marks are deposited in the nucleus and are proposed to function in many nuclear regulatory processes, including microRNA and messenger RNA processing. Despite the widespread use of these pathways, the underlying regulatory programs that influence m^6^A deposition patterns across the transcriptome are poorly characterized. The TARBP2-mediated pathway described here adds a regulatory dimension to RNA methylation and its crucial role in targeted RNA turnover in the nucleus. Importantly, we have discovered that the increased activity of this pathway also promotes lung cancer growth. Employing a network analytical approach, we have identified and functionally validated key factors that lie upstream and downstream of TARBP2 that take part in its oncogenic role in lung cancer. The importance of this TARBP2-mediated regulatory program in multiple cancer types further highlights its central role as a key regulator of gene expression.

## RESULTS

### TARBP2 binding results in increased intron retention and destabilization in the nucleus

Initially, to characterize the regulatory consequences of TARBP2 modulation, we performed siRNA-mediated knockdown of TARBP2 followed by high-throughput RNA sequencing. We then asked whether TARBP2-bound transcripts, defined by analysis of TARBP2 HITS-CLIP data (Goodarzi et al., 2014), show a concerted change in abundance. Consistent with our previous findings obtained from microarrays, we observed that transcripts directly bound by TARBP2 were significantly upregulated when TARBP2 was silenced (Figure 1A). However, the molecular mechanisms linking TARBP2 binding to transcript destabilization were unknown. A key to understanding this mechanism was the observation that, in our previously published TARBP2 HITS-CLIP data (Goodarzi et al., 2014), TARBP2 shows pervasive binding to intronic sequences (e.g. Figure S1A). This suggested that TARBP2 binds to pre-mRNAs and may function in the nucleus by influencing RNA processing and clearance. The major known pathway for RNA degradation in the nucleus involves the targeted destruction of incorrectly spliced transcripts by the RNA surveillance machinery (Kilchert et al., 2016). We hypothesized that TARBP2 may take advantage of this pathway by inhibiting efficient processing of its bound introns, resulting in nuclear retention and degradation of its target transcripts. To test the response of TARBP2-bound introns upon modulation of TARBP2 levels, we used high-throughput transcriptomic profiling measurements from control and TARBP2 knockdown cells to assess the changes in abundance of TARBP2 target transcripts at the exonic and intronic levels.

**Figure 1:**
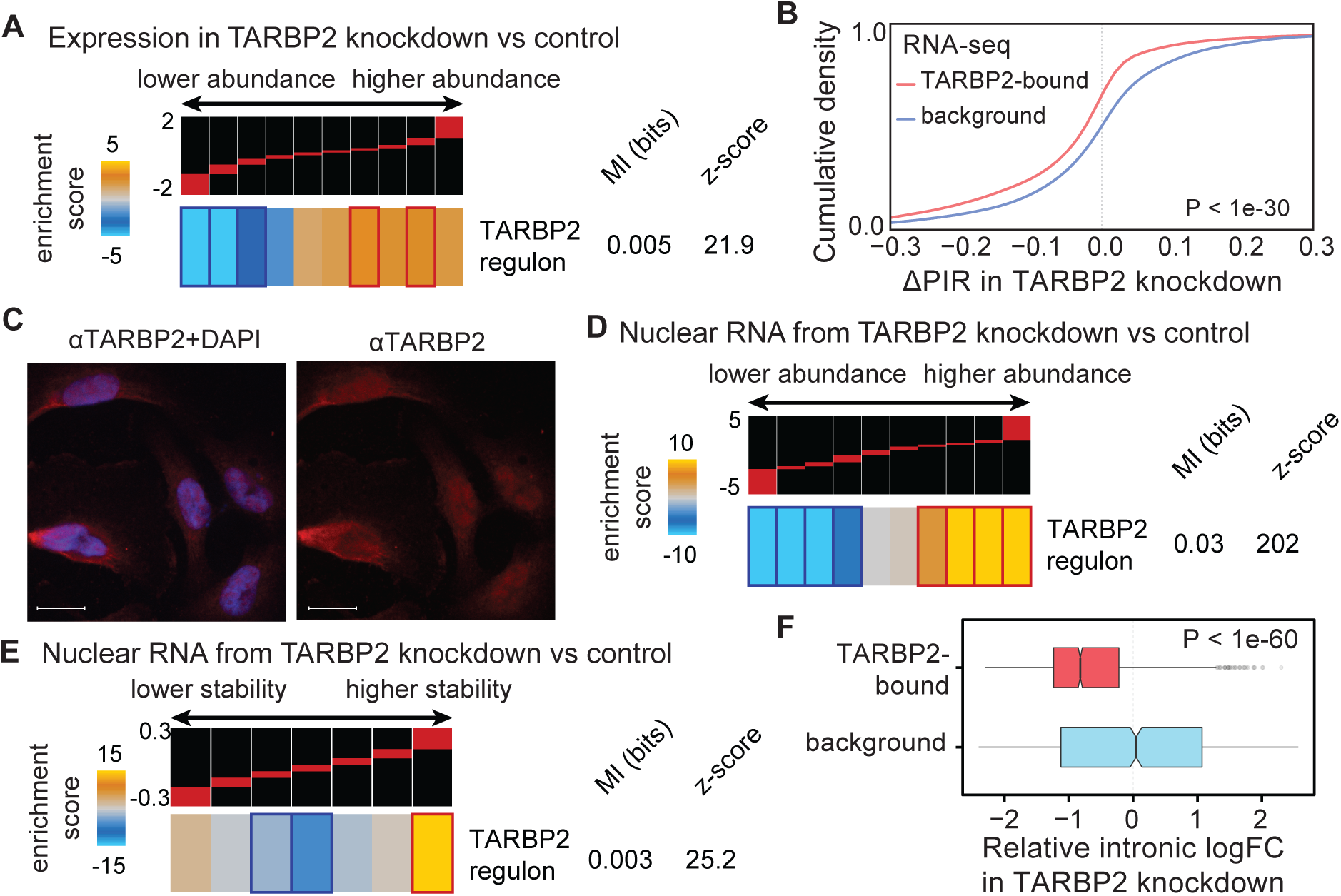
Nuclear TARBP2 binding increases pre-mRNA intron retention and decreases transcript stability. **(A)** A heatmap showing the enrichment of the TARBP2 regulon among transcripts with higher expression in TARBP2 knockdown compared to control MDA-LM2 breast cancer cells. Genes are sorted based on their expression changes in TARBP2 knockdown cells, from down-regulated (left) to up-regulated (right), and grouped into equally populated bins that are visualized as columns. The red bar on top of every column shows the range of log-fold change values for the genes in its corresponding bin. In the heatmap, high enrichment scores are represented by gold, and correspond to bins with enrichment of TARBP2-bound transcripts, while blue represents depletion of TARBP2-bound transcripts. Statistically significant enrichments and depletions, based on hypergeometric tests, are marked with red and dark-blue borders, respectively. Also included are mutual information (MI) values and their associated z-scores (see Methods). **(B)** Cumulative distribution of changes in percent intron retention (PIR) for TARBP2-bound introns in TARBP2 knockdown relative to control MDA-LM2 breast cancer cells (red line). A background set containing a similar number of introns with no evidence of TARBP2 binding (based on HITS-CLIP) is included as a control (blue line). *P* calculated using a Mann-Whitney *U* test. **(C)** Immunofluorescence staining for TARBP2 (red) and staining with DAPI (blue). Shown are z-slices obtained from confocal microscopy imaging of MDA-LM2 cells. Scale bars, 25µm. **(D)** Enrichment of the TARBP2 regulon among the transcripts upregulated in nuclear RNA from TARBP2 knockdown compared to control MDA-LM2 breast cancer cells. **(E)** Enrichment of the TARBP2 regulon among transcripts that are stabilized in the nucleus upon TARBP2 knockdown in MDA-LM2 cells. **(F)** Relative fold-change of percent intron retention for TARBP2-bound introns in nuclear RNAs isolated from TARBP2 knockdown and control MDA-LM2 breast cancer cells.

We annotated TARBP2-bound introns and quantified their abundance relative to their flanking exons using a probabilistic model (MISO; Katz et al., 2010). To measure the global impact of TARBP2 silencing on the splicing of TARBP2-bound introns, we quantified the change in percent intron retention (PIR) across all TARBP2-bound introns in TARBP2 knockdown and control cells. As shown in Figure 1B, we observed a significant shift towards an increased rate of intron splicing when TARBP2 is silenced.

Pervasive TARBP2 binding to intronic sequences implies that this dsRBP is present in the nucleus, which is further supported by a previous study that observed tagged TARBP2 in the nucleus of HeLa cells (Laraki et al., 2008). The Human Protein Atlas also provides immunofluorescence staining showing TARBP2 localized to the nucleoplasm of HeLa and MCF7 cells. To further verify that endogenous TARBP2 is present in the nucleus, we performed immunofluorescence staining followed by confocal microscopy to assess the cellular localization of TARBP2. Consistent with our model, TARBP2 was present in the nucleus as well as the cytoplasm of MDA-LM2 cells (Figure 1C). As will be discussed below, this observation was further confirmed through subcellular fractionation followed by label-free mass-spectrometry and western blotting for TARBP2.

The presence of TARBP2 in the nucleus, as well as its binding to intronic sequences, suggested that TARBP2 may mediate the processing and stability of its target transcripts in the nucleus. To investigate this possibility, we performed high-throughput sequencing on nuclear RNA from TARBP2 knockdown and control cells. We observed a highly significant increase in the expression of the TARBP2 regulon in the nucleus (Figure 1D); this effect was similar to but substantially stronger than the effect observed in total RNA (Figure 1A). To confirm that the observed nuclear upregulation of the TARBP2 regulon is due to post-transcriptional modulation of RNA stability, we performed whole-genome transcript stability measurements by using α-amanitin to inhibit RNA polymerase II and gene expression profiling to assess changes in relative transcript stability in TARBP2 knockdown and control cells. Consistent with our hypothesis and our previous observations, we noted a significant enrichment of TARBP2-bound transcripts among those stabilized in TARBP2 knockdown cells (Figure 1E). To further investigate the effect of TARBP2 binding to introns we measured changes in intron retention by analyzing nuclear RNA-seq data from TARBP2 knockdown and control cells. Silencing TARBP2 resulted in a substantial and significant decrease in the abundance of TARBP2-bound introns compared to introns not bound by TARBP2 (Figures 1F, S1B). On average, target transcripts that are upregulated upon TARBP2 silencing showed a 5% reduction in retention of their TARBP2-bound introns (P<1e-100).

### Nuclear TARBP2 interacts with mRNA processing and export factors

In order to identify the molecular components through which nuclear TARBP2 exerts its regulatory effects, we carried out an unbiased search for its interacting protein partners. We performed immunoprecipitation of both nuclear and cytoplasmic TARBP2, along with an IgG control, to identify proteins that interact with TABP2 in the nucleus (Figure 2A). We searched for RNA-binding protein complexes that were significantly overrepresented in the TARBP2 immunoprecipitation samples compared to IgG co-precipitated proteins (StringDB; Szklarczyk et al., 2015). Ranking high on the list of statistically significant complexes were two involved in RNA processing: a complex containing the RNA processing factor WTAP, and another containing the nuclear pore-associated protein TPR (Figure S2A-B). This analysis suggested that TARBP2 may produce its effect on RNA stability through interactions with these proteins and their associated pathways.

**Figure 2:**
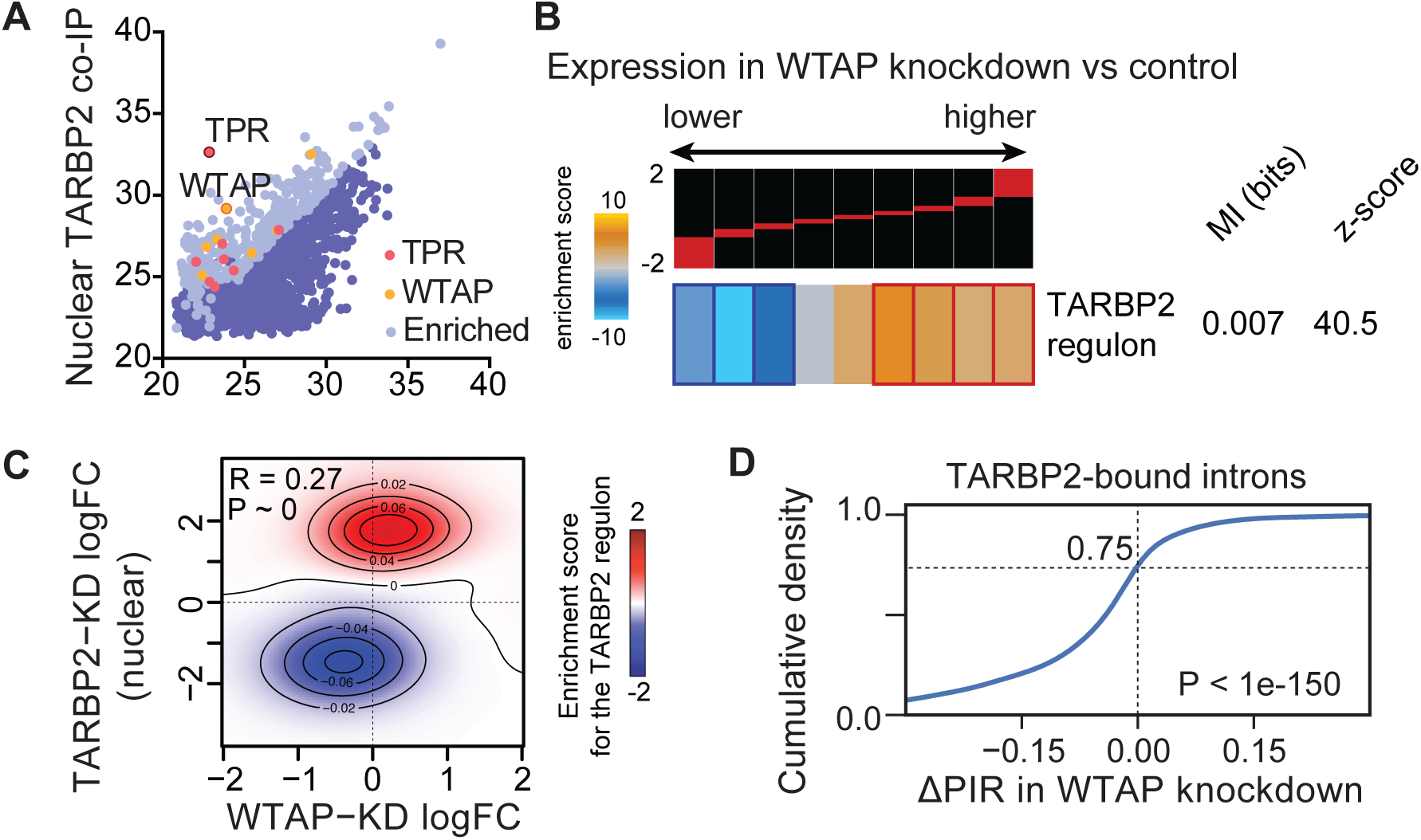
TARBP2 regulates intron retention through interactions with N(6)-methyladenosine methyltransferase associated factors. **(A)** Scatter plot of mass spectrometry data showing proteins that co-immunoprecipitated with TARBP2 versus control IgG in nuclear lysate. Shown are the average of three replicates across all detected proteins. Proteins enriched in the TARBP2 co-IP samples are shown in light blue. In red are TPR and associated proteins, while WTAP and associated proteins are shown in gold. **(B)** Enrichment of the TARBP2 regulon among transcripts that are upregulated in WTAP knockdown compared to control cells (MDA-LM2 background). Also included are the mutual information value (MI) and the associated *z*-score. **(C)** Two-dimensional heatmap showing enrichment of TARBP2-bound transcripts in the group of transcripts that is upregulated in both TARBP2 and WTAP knockdown cells. **(D)** Cumulative distribution of changes in percent intron retention (PIR) in WTAP knockdown relative to control MDA-LM2 breast cancer cells. *P* calculated based on a Wilcoxon signed-rank test.

### TARBP2 recruits m^6^A methylation machinery to mark target intronic sequences

To examine the role of WTAP in modulating expression of the TARBP2 regulon, we carried out siRNA-mediated knockdown of WTAP followed by RNA-seq to quantify both the expression of TARBP2 targets and the processing of their introns. As shown in Figure 2B, silencing WTAP resulted in a significant increase in the expression of transcripts bound by TARBP2. Importantly, these TARBP2-bound transcripts were significantly overrepresented in the set of transcripts upregulated in both TARBP2 and WTAP knockdown cells, and the gene expression changes resulting from TARBP2 and WTAP knockdown are highly correlated (R=0.27; Figure 2C). Moreover, the upregulation of the TARBP2 regulon upon WTAP silencing coincided with an increase in the splicing of TARBP2-bound introns (75% of introns with PIR below zero, *P*∼0; Figure 2D). TARBP2 target transcripts with increased expression in WTAP knockdown cells showed a 9% average reduction in PIR for their TARBP2-bound introns (*P*<1e-100). Furthermore, analysis of a previously published WTAP PAR-CLIP dataset (Liu et al., 2014) revealed a significant overlap between TARBP2-bound introns and WTAP binding sites located in expressed introns and their flanking exons (Figure S2C). We included flanking exons of the TARBP2 bound introns in this analysis as many regulatory elements that influence intron splicing bind exonic regulatory elements. Therefore, WTAP silencing resulted in decreased intron retention and increased expression of the TARBP2 regulon, which further establishes this protein as a component of this TARBP2-mediated pathway.

The observed change in intron retention for TARBP2-bound introns in response to WTAP knockdown is consistent with the known function of WTAP as an RNA processing factor. However, WTAP also serves as the regulatory component of the m^6^A methyltransferase complex (Liu et al., 2014; Ping et al., 2014), and the extent to which these two functions overlap is not known. Thus, the impact of WTAP on the TARBP2 regulon may be dependent on or independent of its role in RNA methylation. To address this question, we first asked whether the interaction between TARBP2 and WTAP has an impact on the methylation status of TARBP2-bound transcripts. We analyzed our previously published nuclear m^6^A co-immunoprecipitation followed by sequencing data (MeRIP-seq; Alarcón et al., 2015a) and we observed a highly significant overlap between introns that are bound by TARBP2 and those that contain an m^6^A mark (Figure 3A). While less than 10% of all expressed introns (and their flanking exons) show evidence of m^6^A methylation, more than half of TARBP2-bound introns contain methylation marks (Figure 3A). These observations suggest a model where TARBP2-mediated recruitment of the methyltransferase complex results in the methylation of its target transcripts. In support of this model, we also observed that METTL3, the enzymatic component of the methyltransferase complex, co-immunoprecipitated with TARBP2, providing further evidence that TARBP2 interacts with the m^6^A methyltransferase complex (Figure 3B). Furthermore, we observed a significant overlap between expressed introns (and their flanking exons) bound by TARBP2 and those bound by METTL3 (PAR-CLIP; Liu et al., 2014) (Figure 3C).

**Figure 3:**
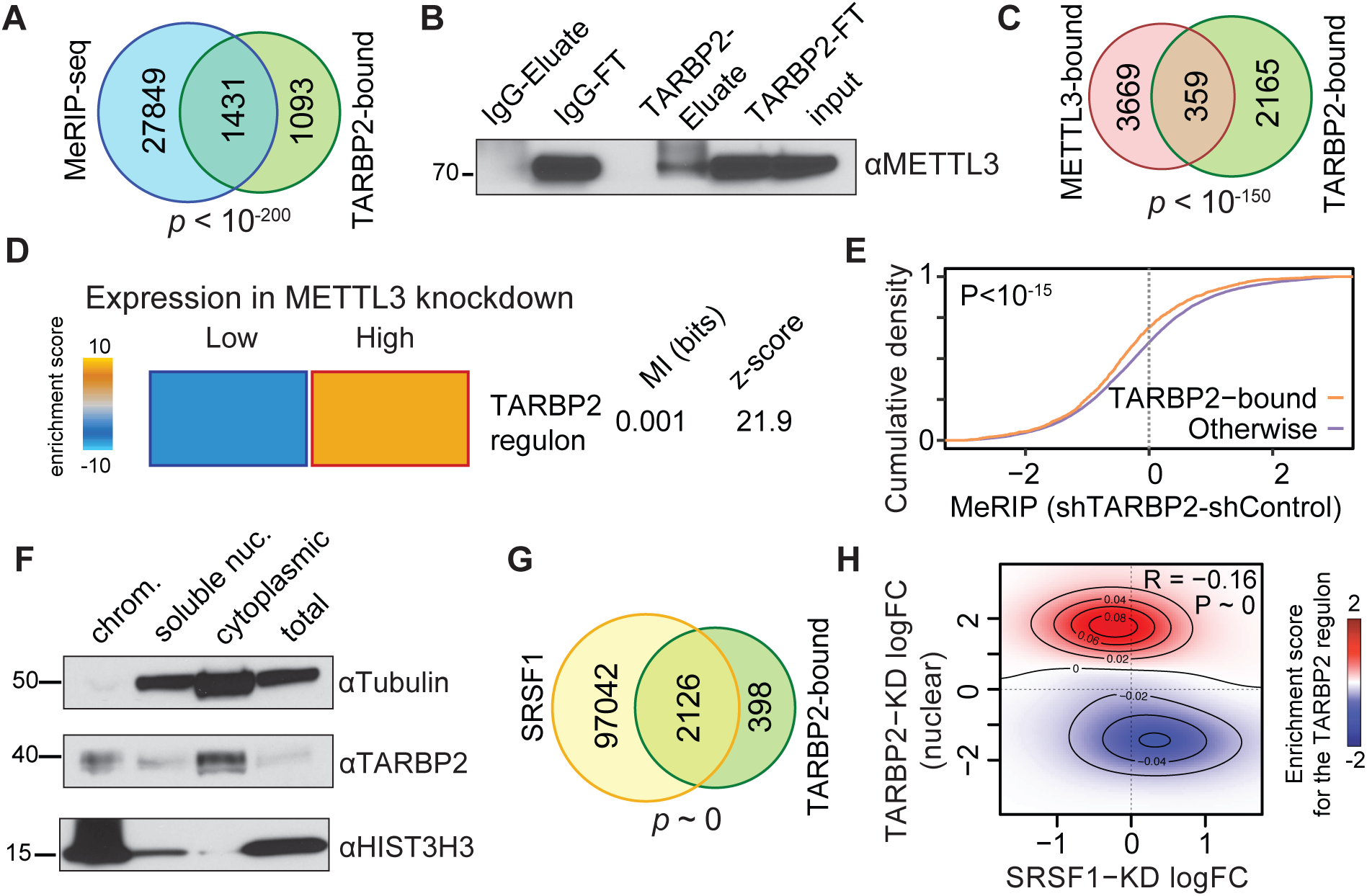
TARBP2-dependent m^6^A modification results in intron retention. **(A)** Venn diagram showing significant overlap between introns and flanking exons bound by TARBP2 and those containing m^6^A marks in MDA-LM2 breast cancer cells. **(B)** Western blot for METTL3 for input, flowthrough and eluate from TARBP2 and IgG immunoprecipitations in MDA-LM2 lysate. **(C)** Venn diagram showing significant overlap between introns and flanking exons bound by TARBP2 and those bound by METTL3 based on a previously published METTL3 PAR-CLIP data (Liu et al., 2014). **(D)** Heatmap showing enrichment of the TARBP2 regulon among transcripts upregulated upon METTL3 knockdown relative to control MDA-LM2 breast cancer cells (data from Alarcón et al., 2015a). The associated mutual information (MI) and *z*-score are shown. **(E)** MeRIP-seq was performed in MDA-LM2 TARBP2 knockdown and control cells. Cumulative distribution graph showing introns bound by TARBP2 have significantly reduced methylation upon TARBP2 knockdown compared to introns not bound by TARBP2. **(F)** Cytoplasmic, soluble nuclear and chromatin-associated protein fractions were collected from MDA-LM2 cells, and western blotting was used to detect TARBP2, tubulin (cytoplasmic) and histone H3 (nuclear). **(G)** Venn diagram showing significant overlap between introns and flanking exons bound by TARBP2 and those bound by SRSF1 (Van Nostrand et al., 2016). **(H)** Two-dimensional heatmap showing enrichment of TARBP2-bound transcripts in the group of transcripts that is upregulated in both TARBP2 knockdown cells and downregulated in SRSF1 knockdown cells.

In order to verify that the regulatory effects of WTAP are mediated through m^6^A RNA methylation, and given that TARBP2 interacts with METTL3, we also analyzed high-throughput RNA-seq data from METTL3 knockdown cells (Alarcón et al., 2015a). Despite the modest reduction in METTL3 levels achieved by knockdown (∼50%), we observed a significant increase in the expression of the TARBP2 regulon (Figure 3D). Finally, to test the causal link between TARBP2 binding and the methylation of its target RNAs, we performed MeRIP-Seq in TARBP2 knockdown and control cells. Consistent with a role for TARBP2 in recruiting the methyltransferase complex, in TARBP2 knockdown cells we observed a significant decrease in the m^6^A signal in the TARBP2-bound introns relative to other expressed introns (Figure 3E).

### TARBP2-dependent m^6^A methylation impacts intron retention and RNA stability

Recently, it was shown that RNA methylation can occur co-transcriptionally, and m^6^A marks can be detected in chromatin-associated RNA (Ke et al., 2017). Analysis of these chromatin-associated RNA m^6^A marks (CA-m^6^A) revealed that there is a significant enrichment of TARBP2-bound introns among chromatin-associated methylated introns (Figure S3A). In contrast, CA-m^6^A marks are largely absent in introns with no evidence of TARBP2 binding (background introns). This suggested that TARBP2 may regulate m^6^A deposition co-transcriptionally. In support of this observation, we blotted for TARBP2 in different sub-cellular compartments, and detected TARBP2 in the chromatin-associated protein fraction, and, to a lesser extent, the soluble nuclear fraction (Figure 3F). This result demonstrates that a portion of nuclear TARBP2 is associated with chromatin, which is consistent with a role for TARBP2 in promoting the co-transcriptional m^6^A methylation of RNA.

To independently confirm the intron retention analysis of our RNA-seq data, we randomly selected a number of TARBP2-bound introns with known methylation sites at nucleotide resolution (CA-m6A and MeRIP-Seq; Alarcón et al., 2015a; Ke et al., 2017). We then used qRT-PCR to measure TARBP2-dependent relative changes in retention of this set of introns using exon-exon and exon-intron spanning primers. In all but one of these cases, we observed a significant decrease in intron retention upon TARBP2 knockdown (Figure S3B). We also used qRT-PCR to confirm that these decreases in intron retention were accompanied by increases in the levels of these mature mRNAs (Figure S3C). As expected from our model, we also noted a significant anti-correlation between changes in intron retention and RNA expression for these transcripts (Figure S3D).

Taken together, our findings support a molecular mechanism of action where TARBP2-mediated methylation of introns interferes with splicing. Based on this model, it is plausible that pre-mRNA m^6^A marks may modulate local binding of regulators of the splicing machinery. To address this possibility, we compiled a list of RNA-binding proteins that differentially bind methylated RNA (Alarcón et al., 2015b; Edupuganti et al., 2017; Liu et al., 2015). We then systematically analyzed the distribution of binding sites of each RBP (derived from CLIP-seq data and known RBP binding motifs, 33 RBPs were included in this analysis) to determine if each candidate RBP binds to introns (and their flanking exons) that are also bound by TARBP2 and/or contain m6A marks. We note that this analysis was limited to proteins with available CLIP-seq data, and therefore does not include all known m^6^A binding proteins. This analysis identified SRSF1 (Serine and arginine rich splicing factor 1), a major modulator of both pre-mRNA splicing and alternative splicing (Das and Krainer, 2014), and a factor that exhibits decreased binding to methylated RNA (Edupuganti et al., 2017), as a top candidate. As shown in Figure 3G, our analysis of SRSF1 CLIP-seq data (Van Nostrand et al., 2016) revealed a highly significant overlap between SRSF1-bound and TARBP2-bound targets. Moreover, thousands of m^6^A sites overlap with SRSF1 binding sites within the TARBP2-bound introns (Figure S3E). These analyses suggest a model where SRSF1-dependent intron splicing is inhibited by TARBP2-dependent m^6^A methylation of introns. To evaluate this hypothesis, we performed RNA-seq on SRSF1 knockdown and control cells, and, consistent with our model, we observed a significant decrease in the expression of the TARBP2 regulon upon SRSF1 knockdown (Figure S3F). Importantly, we observed that gene expression changes induced by TARBP2 and SRSF1 knockdown were significantly anti-correlated (Figure 3H). Moreover, our analysis shows that transcripts bound by TARBP2 were significantly overrepresented in the set of transcripts that was both upregulated in TARBP2 knockdown cells and downregulated in SRSF1 knockdown cells (Figure 3H).

Based on our analyses, in addition to SRSF1, other regulators of RNA splicing that are known to differentially bind methylated RNA may also contribute to TARBP2-dependent intron retention. One potential additional factor is the splicing regulator HNRNPC, which was recently shown to preferentially bind m^6^A methylated RNA (Liu et al., 2015). We observed a significant overlap between the target introns bound by HNRNPC (Zarnack et al., 2013) and those bound by TARBP2 (Figure S3G). Consistently, we also observed an increase in the expression of the TARBP2 regulon when HNRNPC is silenced (Liu et al., 2015) (Figure S3H). We similarly assessed the role of YTHDC1, a known nuclear m^6^A reader that has been implicated in splicing regulation (Xiao et al., 2016; Xu et al., 2014). However, we observed only a slight overlap between TARBP2 and YTHDC1 binding on introns (Xiao et al., 2016). Consistently, YTHDC1 knockdown resulted in little change in the expression of the TARBP2 regulon (data not shown). Together, our findings implicate SRSF1 as a key splicing regulator that is repelled upon methylation of its binding sites on TARBP2-bound introns. Subsequently, decreased SRSF1 binding results in increased intron retention and decreased expression of the mature transcript. In addition to SRSF1, HNRNPC may play a minor role in this regulatory process.

### TARBP2 delivers its target transcripts to the nuclear RNA surveillance complex for degradation

TPR, a nuclear pore-associated factor, was the highest-ranking TARBP2 interacting protein in the nucleus. Given the known role of this protein in nuclear RNA surveillance and degradation of mis-spliced transcripts (Coyle et al., 2011; Krull et al., 2004; Rajanala and Nandicoori, 2012), its interaction with TARBP2 suggested a direct mechanism for the nuclear retention and degradation of TARBP2-bound transcripts. Consistent with this, the TARBP2 regulon was significantly enriched among transcripts that were upregulated upon TPR knockdown (Figure 4A). Gene expression changes in TARBP2 and TPR knockdown cells were also positively correlated, highlighting the functional overlap of these two proteins (Figure 4B). Furthermore, TARBP2-bound transcripts were significantly enriched among the set of transcripts that was upregulated in both TARBP2 and TPR knockdown cells (Figure 4B). If these regulatory consequences of TPR are mediated through its function as a component of the nuclear RNA surveillance machinery, other factors in this complex should have a similar impact on the TARBP2 regulon. To evaluate the role of the nuclear RNA surveillance machinery in TARBP2-mediated transcript destabilization, we analyzed previously reported iCLIP data for EXOSC10 (Macias et al., 2015), a catalytic component of the nuclear exosome complex. As shown in Figure S4A, TARBP2-bound transcripts are significantly enriched among those that are also bound by EXOSC10, consistent with these transcripts being targeted by the surveillance machinery for degradation. Moreover, an additional independent RNA-seq dataset from cells with EXOSC10 knockdown (Macias et al., 2015) revealed a significant increase in the expression of the TARBP2 regulon (Figure S4B). We confirmed this observation by performing RNA-seq in cells with EXOSC10 or XRN2 (a nuclear 5’ to 3’ exonuclease) knockdown (Figures 4C, S4C). In both these cases, we noted a significant increase in the expression of the TARBP2 regulon, confirming that the destabilization of this regulon is contingent on the catalytic activity of the nuclear RNA decay machinery.

**Figure 4:**
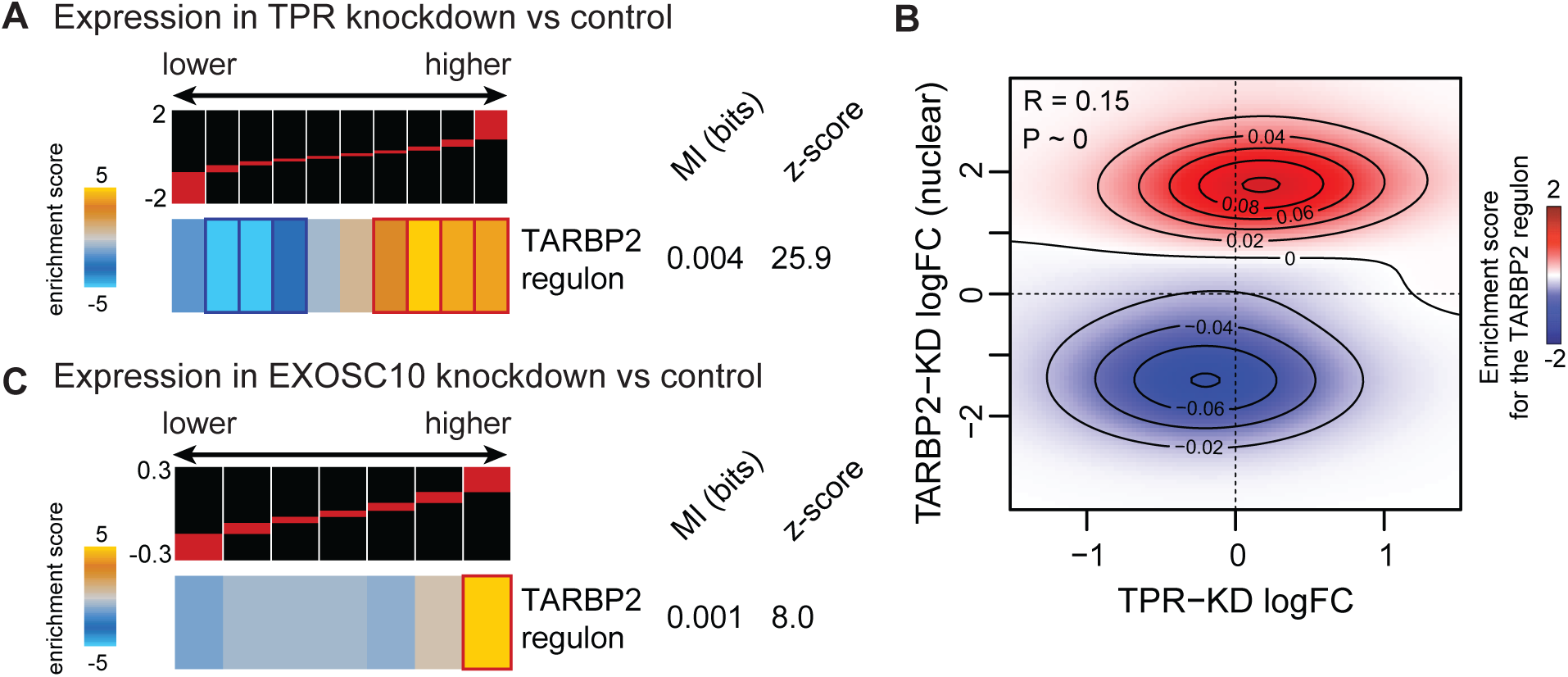
RNA processing and nuclear exosome factors control degradation of the TARBP2 regulon. **(A)** Heatmap showing enrichment of the TARBP2 regulon among genes upregulated in TPR knockdown compared to control MDA-LM2 breast cancer cells. **(B)** Two-dimensional heatmap showing enrichment of TARBP2-bound transcripts in the group of transcripts that is upregulated in both TARBP2 knockdown cells and TPR knockdown cells. Red indicates TARBP2 regulon enrichment, blue depletion. Heatmap showing enrichment of the TARBP2 regulon among transcripts upregulated in EXOSC10 knockdown relative to control MDA-LM2 breast cancer cells.

### Upregulation of the TARBP2 pathway is associated with lung cancer

We have previously described a role for aberrant TARBP2 activity in metastatic breast cancer (Goodarzi et al., 2014). However, the broader role of this pathway in normal cell physiology and human disease remains largely unknown. To address this problem, we performed an unbiased search for evidence of aberrant TARBP2 activity across human cancers profiled in The Cancer Genome Atlas (TCGA) (Cancer Genome Atlas Research Network et al., 2013). Consistent with our previous findings, we observed a significant association between the TARBP2 gene expression signature and breast cancer. However, the TARBP2 signature also showed broad up-regulation in several other cancer types, with the strongest association observed in lung cancer (Figure S5A). This observation was also validated in independent lung cancer datasets, which showed strong upregulation of TARBP2 (Figures 5A, S5B). Importantly, we also observed a strong association between TARBP2 expression and survival in lung cancer patients (Figure 5B). To independently confirm this observation, we used qRT-PCR to measure TARBP2 mRNA levels in lung tumor samples from a cohort of lung cancer patients (including 20 stage I, 14 stage II, and 6 stage III) as well as lung tissue from healthy individuals. As shown in Figure 5C, we observed a substantial and significant upregulation of TARBP2 in lung adenocarcinoma compared to healthy tissue samples.

**Figure 5:**
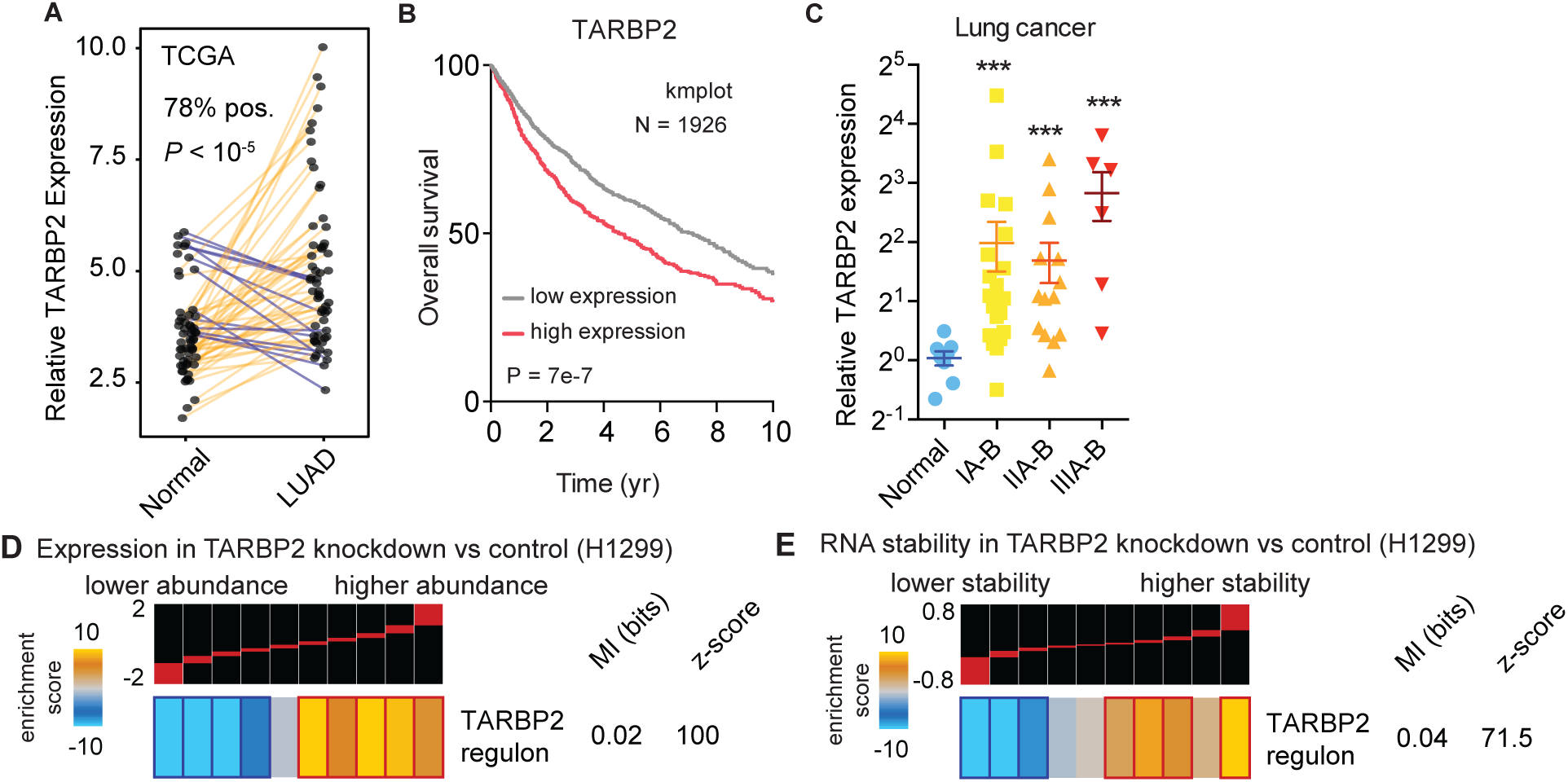
TARBP2 is associated with clinical outcome in lung cancer. **(A)** Upregulation of TARBP2 mRNA in lung adenocarcinoma (LUAD) samples relative to their matched control in The Cancer Genome Atlas dataset. *P* was calculated using Wilcoxon tests. **(B)** Kaplan-Meier survival curve showing overall survival of non-small cell lung cancer patients as a function of TARBP2 expression (Győrffy et al., 2013). *P* calculated using a log-rank test. **(C)** Relative mRNA levels of TARBP2 in normal lung tissue and lung adenocarcinoma stages I-III (N=96) measured using qRT-PCR. *P* calculated using Mann Whitney *U* test. **(D)** Heatmap showing enrichment of the TARBP2 regulon among transcripts upregulated in TARBP2 knockdown H1299 lung cancer cells. The associated mutual information (MI) and z-score are also shown. **(E)** TARBP2-dependent transcript stability was measured by treating TARBP2 knockdown and control H1299 lung cancer cells with α-amanitin and performing RNA-seq. TARBP2-bound transcripts are enriched among transcripts with increased stability in TARBP2 knockdown H1299 lung cancer cells.

Consistent with our observations in breast cancer cell lines, silencing TARBP2 in H1299 lung cancer cells resulted in a significant upregulation and stabilization of the TARBP2 regulon (Figure 5D-E). We also observed this upregulation in the lung cancer cell lines A549 and H1650 (Figure S5C-D). Together, these analyses provide evidence that TARBP2 is strongly associated with lung cancer and that TARBP2-mediated modulation of RNA stability occurs in lung cancer cells.

### TARBP2 promotes lung cancer in *in vivo* models

As our analysis of clinical datasets provides compelling evidence that TARBP2 plays a role in lung cancer, we sought to experimentally test this hypothesis using xenograft mouse models. We injected TARBP2 knockdown and control H1299 lung cancer cells into the venous circulation of immunodeficient mice and then measured cancer cell growth in the lung over time using *in vivo* bioluminescence imaging. While TARBP2 knockdown resulted in only a modest decrease in *in vitro* cell proliferation (Figure S6A), we observed a significant reduction in growth in the lung by cells with TARBP2 knockdown compared to control cells (Figure 6A).

**Figure 6:**
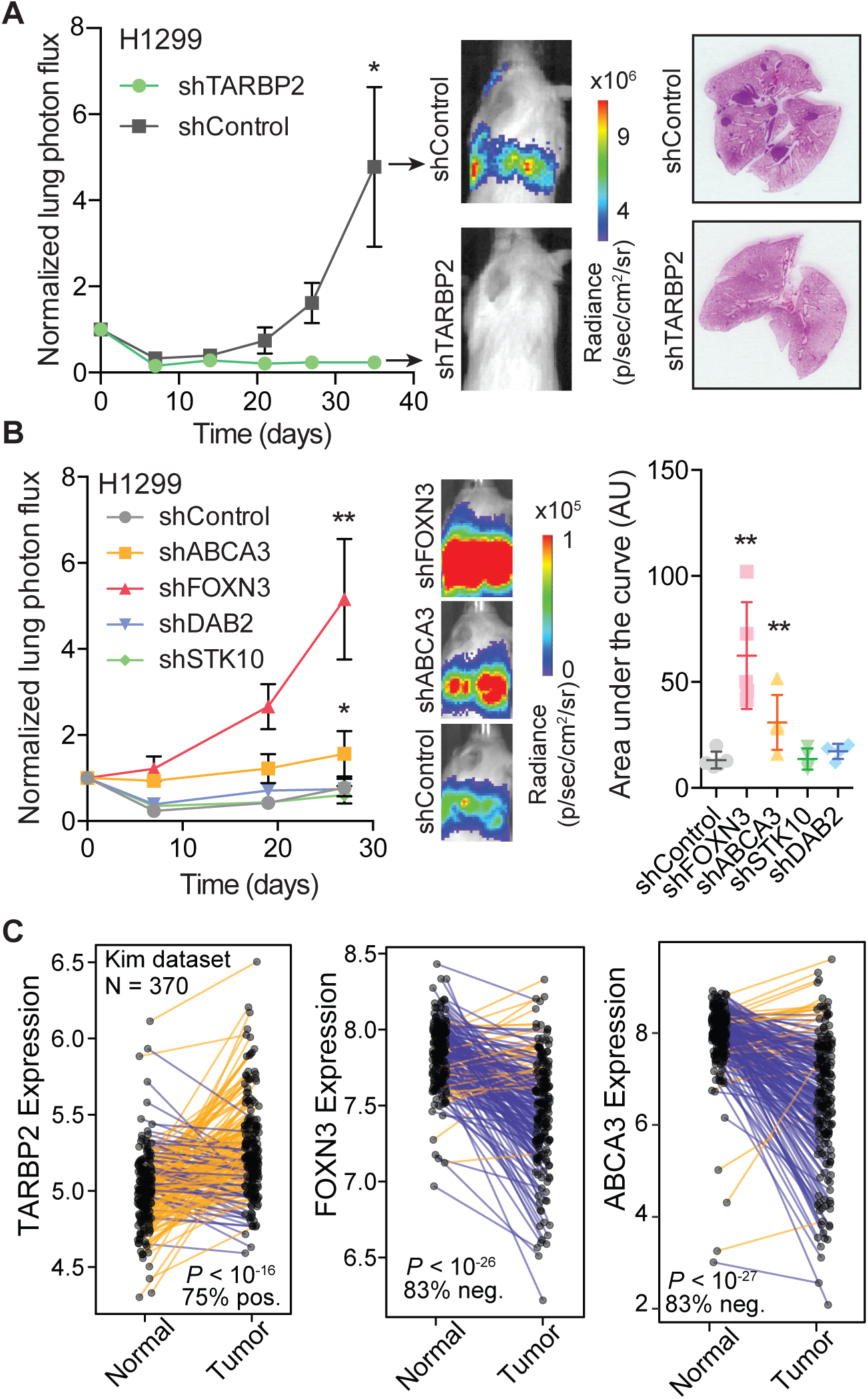
TARBP2 targets ABCA3 and FOXN3 promote lung cancer growth *in vivo*. **(A)** Plots showing lung bioluminescence signal over time in mice injected with H1299 lung cancer cells expressing a TARBP2 targeting shRNA or a control shRNA. Representative H&E stained lungs are also shown (N=5 per cohort). **(B)** Plots showing lung bioluminescence signal over time in mice injected with H1299 lung cancer cells expressing shRNAs targeting ABCA3, FOXN3, DAB2, STK10, or a control shRNA (N=4-5 per cohort). **(C)** Relative TARBP2, ABCA3 and FOXN3 mRNA expression in paired normal and lung cancer samples (Kim et al., 2013). *P*-value was calculated using the Wilcoxon test.

In order to identify TARBP2 targets that act downstream of TARBP2 to impact lung cancer, we searched for transcripts that were directly bound by TARBP2, had TARBP2-dependent decreased expression and stability, and were negatively correlated with TARBP2 expression in clinical lung cancer datasets (Figure S6B). From this list, we selected the four highest ranking targets that also show evidence of methylation and TARBP2-dependent intron retention, namely FOXN3, ABCA3, DAB2, and STK10. We tested the impact of silencing these genes on lung cancer growth in xenograft models by injecting H1299 cells stably expressing an shRNA against each of these genes. As shown in Figure 6B, silencing FOXN3 and ABCA3 significantly increased lung cancer growth, while decreasing STK10 and DAB2 expression had no significant effect. Moreover, this effect was independent of *in vitro* cell proliferation rates, which showed no significant change upon target gene knockdown (Figure S6C). We also performed qRT-PCR to measure the relative levels of the mature and pre-mRNA of these targets, and observed a significant increase in the mature mRNA levels and significant decrease in the relative pre-mRNA levels of ABCA3 and FOXN3 (Figure S6D). To further assess the clinical relevance of these functional target genes in human disease, we performed a set of additional analyses using clinical datasets. We analyzed a dataset of gene expression profiles from a large cohort of matched normal and lung tumor samples collected from patients (Kim et al., 2013), and we confirmed that TARBP2 is also significantly upregulated in this data (Figure 6C). We then assessed the changes in the expression of FOXN3 and ABCA3 in this same dataset, and consistent with their proposed roles as tumor suppressors, we observed a highly significant reduction in their expression in lung cancers, and found that their expression was significantly correlated with that of TARBP2 (Figures 6C, S6E). Together, these data are consistent with a model where TARBP2 decreases expression of FOXN3 and ABCA3, leading to increased lung cancer growth.

### The transcription factor ZNF143 drives the aberrant upregulation of TARBP2

While our findings provide a molecular understanding of how TARBP2 promotes cancer growth and progression, they did not explain how cancer cells achieve TARBP2 overexpression. Analysis of the TCGA-LUAD (lung adenocarcinoma) dataset showed a general association between TARBP2 expression and TARBP2 genomic copy number (Figure S7A). However, copy number variation alone is not sufficient to explain the magnitude of TARBP2 upregulation in lung cancer. In order to identify the regulatory pathway that drives TARBP2 overexpression in lung cancer, we performed a systematic search for known transcription factors that were significantly co-expressed with TARBP2 in multiple independent cancer datasets. We also performed promoter sequence analysis and ChIP-seq data mining (ENCODE) to identify transcription factors that potentially act as upstream regulators of TARBP2. This exercise yielded a list of five potential candidates. To ask if any of these candidates regulate TARBP2 expression, we performed siRNA-mediated knockdown of each gene in H1299 lung cancer cells followed by qRT-PCR (Figure S7B). Of these candidates, only ZNF143 knockdown resulted in a significant reduction in TARBP2 expression, consistent with ZNF143 controlling TARBP2 transcription. This result was observed with independent ZNF143-targeting siRNAs in additional lung and breast cancer cell lines (Figures 7A, S7C). Moreover, ChIP-seq datasets from multiple cell lines show evidence of strong binding of ZNF143 at the TARBP2 promoter, with the ChIP peak in this region containing a close match to the ZNF143 consensus binding site (Figure 7B). Our analysis also revealed a significant correlation between expression of ZNF143 and TARBP2 in lung cancer gene expression data (Figure S7D). Finally, consistent with ZNF143 playing a role in modulating TARBP2 transcription, we found a significant association between ZNF143 expression and survival in lung cancer patients, as well as a significant upregulation of ZNF143 levels in lung cancers compared to matched normal tissue in the TCGA-LUAD (lung adenocarcinoma) dataset (Figures 7C-D). Together, these results provide strong evidence that the transcription factor ZNF143 increases the expression of TARBP2 in breast and lung cancers.

**Figure 7:**
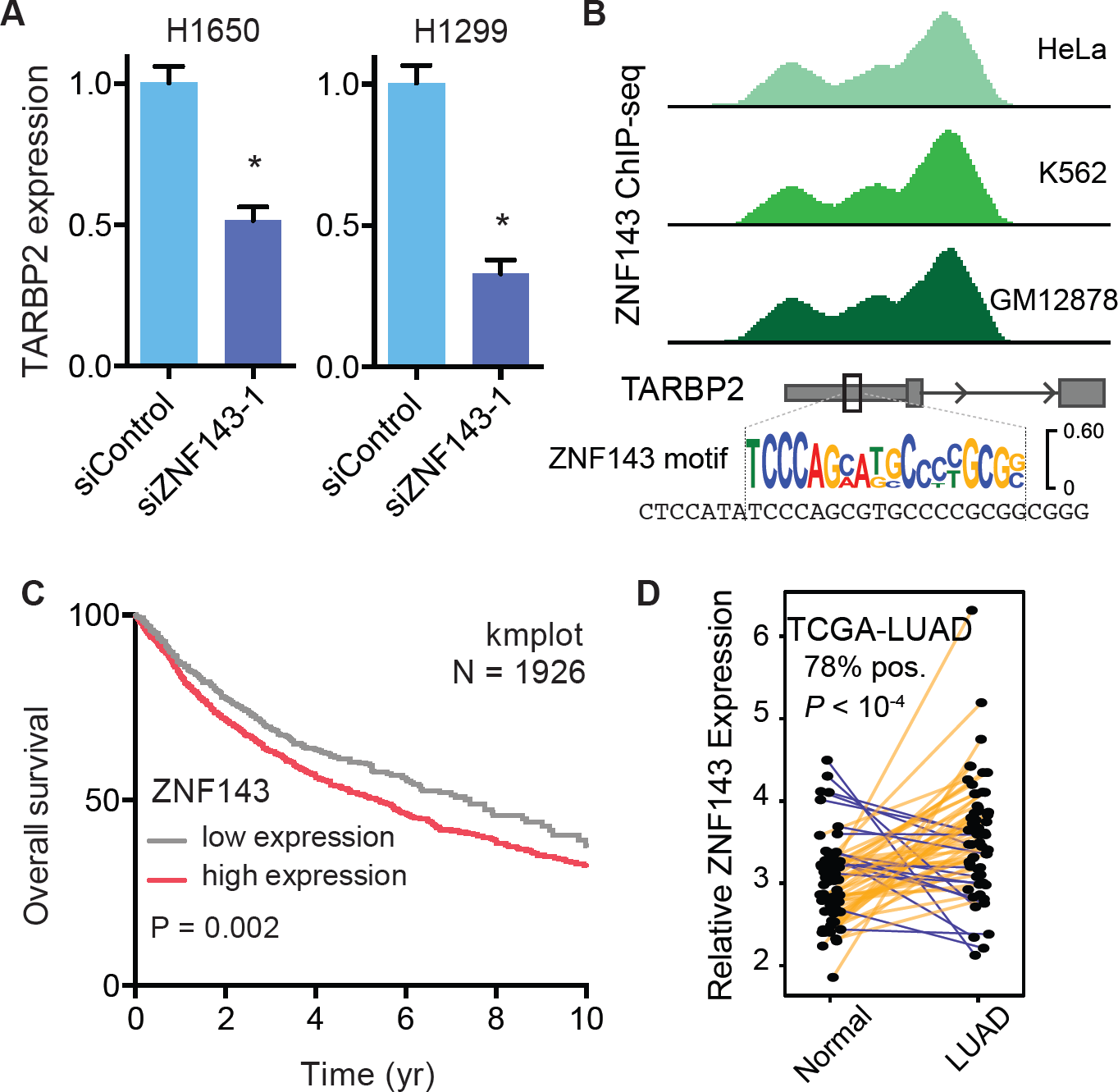
ZNF143 regulates TARBP2 expression in breast and lung cancer. **(A)** Relative TARBP2 mRNA levels were measured by qRT-PCR in H1650 and H1299 lung cancer cells transfected with siRNAs targeting ZNF143 or a control siRNA (N=3). **(B)** ENCODE ChIP-seq tracks from three different cell lines that show evidence of ZNF143 binding to the promoter region of TARBP2. These ChIP peaks are also located at a strong sequence match to the ZNF143 consensus motif. **(C)** Kaplan-Meier survival curve showing overall survival of non-small cell lung cancer patients as a function of ZNF143 expression (Győrffy et al., 2013). *P* was calculated using a log-rank test. **(D)** Relative ZNF143 mRNA expression in paired normal and tumor samples from TCGA-LUAD dataset (*P* based on a Wilcoxon test).

## DISCUSSION

Here, we describe a novel oncogenic post-transcriptional regulatory program controlled by the double stranded RNA binding protein TARBP2. First described as a protein that binds the HIV TAR element, TARBP2 also has a role in miRNA processing (Chendrimada et al., 2005; Gatignol et al., 1991; Kim et al., 2014). Our previous findings demonstrated that TARBP2 regulates RNA stability through the direct binding of RNA structural elements on hundreds of transcripts (Goodarzi et al., 2014). In this study, we dissect the molecular mechanisms through which TARBP2 controls transcript stability, and show that TARBP2 directly controls the stability of its bound targets via co-transcriptional recruitment of the METTL3 methyltransferase complex, resulting in intron methylation and subsequent retention of the intron followed by degradation of the transcript by the nuclear exosome.

Intron retention is a well-documented mechanism of regulating RNA stability. Here, we establish a novel link between TARBP2 intronic binding, m^6^A methylation, and controlled intron retention leading to transcript degradation in the nucleus. Although RNA methylation marks impact a variety of developmental and disease processes (Zhang et al., 2017a, 2017b; Zhao et al., 2017), their mechanistic effects have not been fully explored. One known mechanism linking m^6^A methylation of transcripts to their stability is a cytoplasmic process mediated by the m^6^A binding protein YTHDF2, in which m^6^A-containing transcripts are bound by YTHDF2 in the cytoplasm, which then recruits the CCR4-NOT complex to accelerate their degradation (Du et al., 2016). The pathway we describe here is distinct in that it occurs in the nucleus, implying that it may act on different sets of transcripts. Interestingly, in *S. pombe*, Mmi1, a YTH domain-containing protein binds specific introns, resulting in their retention and subsequent targeted nuclear degradation by the exosome. Although methylation of the target introns has not been demonstrated, this suggests existence of a more general link between intron methylation, retention, and nuclear decay (Kilchert et al., 2015). Building on this mechanism, we also found that a fraction of TARBP2 is associated with chromatin, and therefore it is plausible that TARBP2 promotes intron methylation co-transcriptionally, consistent with a recent report that m^6^A marks are deposited on nascent pre-mRNA (Ke et al., 2017). We speculate that the mode of TARBP2-dependent post-transcriptional regulation we describe here may be advantageous because it allows for the fast decoupling of expression of the TARBP2 regulon from transcriptionally controlled levels. This may be beneficial for promoting oncogenesis and metastasis as it could allow for rapid adaptation to changes in the microenvironment.

Interestingly, although we found a highly significant association between TARBP2 intron binding and RNA methylation, the methylation sites do not necessarily overlap with TARBP2 binding sites. This suggested that additional factors interact with the methylated sites to promote intron retention. By analyzing our data, along with publically available datasets, we found strong associations between TARBP2-mediated intron retention and the splicing regulator proteins SRSF1 and HNRNPC. Our results are consistent with a mechanism where methylation interferes with the ability of SRSF1 to bind and promote intron processing, and potentially enhances the ability of HNRNPC to bind and inhibit intron processing. HNRNPC has been shown to compete with U2AF65 (Zarnack et al., 2013), a function that could inhibit spliceosome assembly, leading to local intron retention.

Transcripts with retained introns may be excluded from cytoplasmic export by TPR, a protein component of the nuclear basket that has a known role in impeding the export of intron containing RNAs (Coyle et al., 2011; Rajanala and Nandicoori, 2012). In this study, we found a physical interaction between TARBP2 and TPR. This TARBP2-TPR interaction suggests that TARBP2 is in close proximity to the nuclear basket, and therefore it is possible that TARBP2-bound intron-containing transcripts are blocked from cytoplasmic export by proximity to TPR. We also identified EXOSC10 and XRN2, catalytic factors of the canonical nuclear decay machinery, as factors responsible for the nuclear degradation of the TARBP2 regulon.

Furthermore, we provide evidence for a functional role for the TARBP2-mediated RNA decay pathway in lung cancer. Intriguingly, we had previously observed that TARBP2 promotes metastatic colonization of the lung by breast cancer cells. Here, we find that the TARBP2 signature is enriched in clinical lung cancer gene expression datasets, and we demonstrate that TARBP2 enhances lung cancer growth *in vivo*, suggesting that the gene expression pattern controlled by TARBP2 is highly suited for promoting oncogenic growth in the lung microenvironment. Also consistent with our results, a previous study reported that knockdown of TARBP2 in H1299 lung cancer cells reduced cell invasion and migration (Shi et al., 2016). Furthermore, our analyses show a robust association between TARBP2 expression and clinical outcome in lung cancer. These findings are consistent with the view that gene expression programs that promote primary tumor growth may also be critical in promoting metastatic colonization of that same organ.

We have also found that in lung cancer, TARBP2 downregulates ABCA3 and FOXN3 expression, leading to increased cancer growth in the lung. ABCA3 is an ATP binding cassette lipid transport protein that is necessary for normal secretion of lung surfactant (Shulenin et al., 2004). Deletion of ABCA3 in mouse genetic models leads to lung tissue injury, inflammation and subsequent proliferation of new cells from progenitors (Rindler et al., 2017). It is possible that these inflammatory and proliferative processes could promote lung cancer growth. It is also possible that ABCA3 directly suppresses cancer cell growth through modulating lipid transport. FOXN3, a second functional TARBP2 target, is a transcriptional repressor that also acts as a cell cycle checkpoint regulator (Pati et al., 1997; Scott and Plon, 2005). Therefore, it is plausible that knockdown of FOXN3 promotes uncontrolled cell division leading to oncogenesis. Although we did not observe a significant change in the *in vitro* proliferation rate of FOXN3 knockdown cells, it is possible that lung-specific microenvironmental cues are required for this effect. Consistent with our findings in lung cancer, downregulation of FOXN3 has been reported to promote proliferation of liver and colon cancer cells (Dai et al., 2016; Sun et al., 2016). Intriguingly, it has been reported that a G to A single-nucleotide polymorphism in the first intron of FOXN3 results in higher FOXN3 expression—it is conceivable that this is a result of disruption of the TARBP2-mediated intron retention pathway (Karanth et al., 2016).

Finally, we have identified ZNF143 as an upstream regulator of TARBP2 expression. Consistent with this, a previous study found an association between high ZNF143 protein levels and poor survival in lung adenocarcinoma (Kawatsu et al., 2014). Intriguingly, a study has identified novel small molecules that inhibit ZNF143 activity (Haibara et al., 2017), pointing towards a possible avenue for inhibition of TARBP2 pro-oncogenic activity.

Taken together, our study reveals a previously unknown post-transcriptional regulatory program that establishes a functional link between the RNA methylation machinery, regulators of RNA splicing, and components of the nuclear RNA surveillance complex. Together, these processes build a novel regulatory mechanism, orchestrated by the RNA-binding protein TARBP2, that modulates the expression of a large set of transcripts in the nucleus. Linking this pathway to both lung and breast cancer progression emphasizes its importance in shaping the gene expression landscape of the cell. However, the same mechanisms might also be employed by other post-transcriptional regulators to modulate expression of their associated regulons. As such, a broader understanding of controlled intron retention and its underlying molecular mechanisms is a crucial step towards achieving a more detailed view of post-transcriptional regulation, as well as exposing new vulnerabilities that can exploited to counter human disease.

## METHODS

### Cell culture

All cells were cultured in a 37°C 5% CO2 humidified incubator. The MDA-MB-231 breast cancer cell line, and its highly metastatic derivative, MDA-LM2, was propagated in DMEM media supplemented with glucose (4.5g/L), 10% FBS, 4mM L-glutamine, 1mM sodium pyruvate, penicillin (100 units/mL), streptomycin (100 µg/mL) and amphotericin (1µg/mL) (Life Technologies). The A549, H1650, and H1299 lung cancer cells were cultured per ATCC recommendations plus penicillin (100 units/mL), streptomycin (100 µg/mL) and amphotericin (1µg/mL).

### shRNA and siRNA-mediated knockdown

DNA transfections were performed using lipofectamine 2000 per the manufacturer’s protocol (Life Technologies). For stable knockdown of target genes by shRNA, pLKO.1 containing shRNAs against the targeted gene was packaged using the ViraSafe lentiviral packaging system (Cell Biolabs).

### RNA Isolation

Total RNA for RNA-seq and quantitative RT-PCR assays was isolated using the Norgen Biotek total RNA isolation kit with on-column DNase treatment per the manufacturer’s protocol.

### Quantitative RT-PCR

Transcript levels were measured using quantitative RT-PCR by first reverse transcribing total RNA to cDNA (SuperScript III, Invitrogen) followed by SYBR Green quantification (Life Technologies) per the manufacturer’s instructions. We used the following primers for qRT-PCR:

### Metastatic colonization assays

Seven-to eight-week-old age-matched female NOD/SCID gamma mice (Jackson Labs) were used for lung metastasis assays. In all cases, 5×10^4^ cells in 100 µL PBS were injected via tail-vein along with cells expressing a neutral hairpin as control. In every case 4-5 mice were included in each cohort. The metastatic growth was tested using both two-way ANOVA as a function of time and sub-line identity and also *t*-test based comparison of area under the curve for each mouse.

### Histology

For gross macroscopic metastatic nodule visualization, mice lungs (from each cohort) were extracted at specific time-points post-injection and 5µm thick lung tissue sections were hematoxylin and eosin (H&E) stained. The number of macroscopic nodules was then recorded for each section. Unpaired *t*-test was used to test for significant variations.

### Cancer cell proliferation

Roughly 10,000 cells were seeded into three 6-well wells and subsequently were trypsinized and viable cells were counted using a hemocytometer at day 1, day 3 and day 5. An exponential model was then used to fit a growth rate for each sample (ln(*N*_*t*-1_/*N*_*1*_)= *rt* where *t* is measured in days). The experiment was performed in biological quadruplicates and unpaired *t*-test was used to test for significant variations.

### Immunofluorescence

MDA-LM2 cells were seeded and incubated for 48 hours in chamber slides, then fixed with 4% paraformaldehyde, washed 3x with PBST (0.1% Tween-20), and blocked for one hour in blocking buffer (5% goat serum, 0.2% fish skin gelatin, 0.2% tween-20). TARBP2 primary antibody (Proteintech 15753) was diluted 1:50 in blocking buffer and incubated with the cells at 4°C overnight. Cells were washed three times with PBST before incubation with anti-rabbit Cy3 secondary (1:1000) (Jackson #715-165-152) in blocking buffer at 37°C for one hour. Cells were washed three times with PBST, the second wash containing 2.5 µg DAPI. Slides were mounted with ProLong Gold antifade reagent (Life Technologies), and images acquired with a Nikon Ti spinning disk confocal microscope at the UCSF Nikon Imaging Center.

### Stability measurements

MDA-LM2 TARBP2 knockdown and control cells were treated with 10 µg/mL α-amanitin. After nine hours, nuclear RNA was harvested from the cells using a Cytoplasmic and Nuclear RNA Purification Kit (Norgen). This RNA was prepared for microarray using TargetAmp-Nano Labeling Kit for Illumina (Epicenter). Labeled RNA was purified using RNeasy Minelute Kit (Qiagen) and submitted for analysis to the Rockefeller University genomics core facility using Illumina HT-12 v4 Expression BeadChip microarrays. The Lumi package in R was used to transform and normalize signal intensities.

### Co-immunoprecipitation and mass spectrometry

MDA-LM2 cells (in biological replicates) were collected by scraping, then centrifuged to pellet. The pellets were then resuspended in buffer LB1 (50 mM HEPES-KOH pH 7.5, 140 mM NaCl, 1 mM EDTA, 10% glycerol, 0.5% Triton X-100 and protease inhibitors), incubated on ice for 15 minutes, and spun for 10 minutes at 2000 rpm. The supernatant was collected (cytosolic fraction) and the nuclei were resuspended in M-PER (Thermo Fisher) plus protease inhibitors, incubated on ice for 10 minutes and spun for 5 minutes. Immunoprecipitation of TARBP2 was carried out by first removing glycerol from anti-TARBP2 and rabbit IgG using an antibody purification kit (Protein A; Abcam) and then conjugating the antibodies to epoxy magnetic beads using the dynabeads antibody coupling kit (Thermo Fisher), all according to the manufacturer’s protocol. The cytosolic and nuclear lysates were then incubated with antibody-conjugated beads for three hours at 4°C with end-over-end rotation. The beads were then washed three times with PBS supplemented with 150mM NaCl and then three times with PBS.

Following immunoprecipitation, proteins were eluted from the antibody-bead conjugate by denaturation in 50µL 8M urea/0.1M ammonium bicarbonate for 30 minutes. Supernatant was removed following reduction (10 mM DTT) and alkylation (40 mM iodoacetamide). Proteins were digested with Endoproteinase LysC (Wako Chemicals) after dilution to 4M urea followed by trypsination (Promega) in 2M urea. The digestion was quenched with 5% formic acid (final concentration) and resulting peptide mixtures were desalted using in-house made C18 Empore (3M) StAGE tips (Rappsilber et al., 2007). Samples were dried and resolubilized in 2% acetonitrile and 2% formic acid and analyzed by reversed phase nano-LC-MS/MS (Ultimate 3000 coupled to a QExactive Plus, Thermo Scientific). After loading on a C18 PepMap trap column (5 µm particles, 100µm x 2 cm, Thermo Scientific) at a flow rate of 3 µl/min, peptides were separated using a 12 cm x 75µm C18 column (3 µm particles, Nikkyo Technos Co., Ltd. Japan) at a flow rate of 200 nL/min, with a gradient increasing from 5% Buffer B (0.1% formic acid in acetonitrile) / 95% Buffer A (0.1% formic acid) to 40% Buffer B / 60% Buffer A, over 75 minutes. All LC-MS/MS experiments were performed in data dependent mode with lock mass of m/z 445.12003. Precursor mass spectra were recorded in a 300-1400 m/z range at 70,000 resolution, and fragment ions at 17,500 resolution (lowest mass: m/z 100) in profile mode. Up to twenty precursors per cycle were selected for fragmentation and dynamic exclusion was set to 60 seconds. Normalized collision energy was set to 27.

Data were searched against a Uniprot human database (July 2014) using MaxQuant (version 1.5.0.30) software and Andromeda search engine (Cox et al., 2014). Oxidation of methionine and N-terminal protein acetylation were allowed as a variable, and cysteine carbamidomethylation as a fixed modification. An ion mass tolerance was set at 4.5 ppm for precursor and 20 ppm for fragment ions. Two missed cleavages were allowed for specific tryptic search. The “match between runs” option was enabled. False discovery rates for proteins and peptides were set to 1%. Protein abundances were represented by LFQ (label free quantitation) values. Data were filtered to exclude contaminants, and reverse database hits. LFQ values were log2(x) transformed and further used for t-test (Tyanova et al., 2016).

### Co-immunoprecipitation of METTL3

Co-immunoprecipitation of TARBP2 and METTL3 was performed using lysate prepared from MDA-LM2 cells. Cells were collected by scraping, then centrifuged to pellet. Pellet was resuspended in M-PER (Thermo Fisher) plus protease inhibitors and incubated on ice 10 minutes. Lysate was clarified by centrifugation for 15 minutes at 20,000 x g at 4°C. TARBP2- or IgG-conjugated dynabeads (prepared as described above) were added to clarified lysate and incubated for two hours at 4°C with end-over-end rotation. Beads were then washed once with PBS supplemented with 150mM NaCl, then washed three times with PBS. Proteins were eluted by resuspending beads in NuPAGE LDS loading buffer and incubating for 10 minutes at 70°C.

### MeRIP-seq

Control and TARBP2 knockdown cells (MDA-LM2 background) were collected and processed as previously described (Alarcon et al., 2015) in biological replicates. 10^7^ cells per sample were lysed using LB1 buffer. The nuclear fraction was then lysed with M-PER buffer (Thermo Scientific) and diluted tenfold in dilution buffer (50 mM Tris-Cl, pH 7.4, 100 mM NaCl) before the immunoprecipitation. Rabbit anti-m6A antibody (Synaptic Systems) and rabbit IgG control bound to protein A Dynabeads (Invitrogen) were used for the immunoprecipitations. The immunoprecipitated RNA was eluted with N6-methyladenosine (Sigma-Aldrich), ethanol precipitated and resuspended in water. RNA was barcoded using ScriptSeq V2 kit (Epicentre) and sequenced at Rockefeller University Genomics Core.

### Western blotting

Cell lysates were prepared by lysing cells in ice-cold RIPA buffer containing 1X protease inhibitors (Roche). Lysate was cleared by centrifugation at 20,000 x g for 15 min at 4°C. Samples were boiled in 1X LDS loading buffer and reducing agent (Invitrogen), separated by running SDS-PAGE, transferred to PVDF (Millipore), blocked using 5% milk and probed using target-specific antibodies. Bound antibodies were detected using horseradish peroxidase– conjugated secondary antibodies, ECL Western Blotting Substrate (Pierce), according to the manufacturer’s instructions. Antibodies: beta-tubulin (Proteintech 66508-1-Ig), TARBP2 (Proteintech 15753), histone H3 (Proteintech 17168-1-AP), METTL3 (Abcam ab195352), TPR (Bethyl A300-827A).

### Subcellular fractionation

Subcellular fractionation was performed using a low-salt method, based on (Méndez and Stillman, 2000). 1.5×10^7^ MDA-LM2 cells were washed in PBS, collected by scraping, and the cell pellet was resuspended in 200µl cold buffer A (10mM HEPES-KOH pH 7.9, 1.5mM MgCl2, 10mM KCl, 340mM sucrose, 10% glycerol, 0.1% Triton X-100, 1mM DTT, 1X halt protease inhibitor cocktail (Pierce)), and rotated end-over-end 10 minutes at 4°C. Sample was then centrifuged at 1,300 x g 5 minutes at 4°C, and supernatant was removed and clarified by centrifuging at 20,000 x g 10 minutes at 4°C. This supernatant was used as the cytoplasmic fraction. The nuclei were washed with 200µl buffer A without Triton X-100, then centrifuged at 1,300 x g 10 minutes 4°C. Nuclei were then resuspended in 100µl buffer B (3mM EDTA, 0.2mM EGTA, 1mM DTT, 1x halt protease inhibitor cocktail (Pierce)), and rotated end-over-end 30 minutes at 4°C. Sample was then centrifuged 1,700 x g 10 minutes at 4°C, and the resulting supernatant was used as the soluble nuclear fraction. The remaining pellet was washed with 200 µl cold phosphate buffered saline, then centrifuged at 1,700 x g 5 minutes at 4°C, and the supernatant discarded. The chromatin pellet was then DNase treated by adding 44µl H2O, 5µl 10X Turbo DNase buffer, and 2 units of turbo DNase (Ambion). The mixture was incubated 10 minutes at 37°C, and was used as the chromatin fraction. Total lysate was prepared by lysing pelleted cells in RIPA (25mM Tris-HCl pH 7.6, 0.15M NaCl, 1% IGEPAL CA-630, 1% sodium deoxycholate, 0.1% SDS) and centrifuging at 20,000 x g 10 minutes to clarify lysate. For Western blotting, 1% of total, and 10% each of the cytoplasmic, soluble nuclear, and chromatin fractions separated by PAGE and detected by immunoblotting as described above.

### RNA-seq library preparation

Unless otherwise specified below, RNA sequencing libraries were prepared using the ScriptSeq-v2 (Illumina) and sequenced on an Illumina HiSeq2000 instrument. RNA-seq libraries for expression profiling of MDA-LM2 cells with shRNA-mediated XRN2 or EXOSC10 knockdown were generated using the QuantSeq 3’ mRNA-Seq library prep kit fwd (Lexogen) per the manufacturer’s protocol, and sequenced on an Illumina HiSeq4000 at UCSF CAT.

## AUTHOR CONTRIBUTIONS

Conceptualization, L.F. and H.G.; Methodology, L.F. and H.G; Formal Analysis, H.S.N. and H.G; Investigation L.F., H.C.B.N., S.Z., M.H., B.D.D., H.M., C.A. and H.G.; Writing – Original Draft, L.F. and H.G.; Writing – Review & Editing, L.F. and H.G.; Supervision, H.G.

## ACKNOWLEDGEMENTS

We are grateful to Sohail Tavazoie and Saeed Tavazoie for reading the earlier versions of this manuscript. We acknowledge the UCSF Center for Advanced Technology (CAT) and the Rockefeller Genomics Resource Center for high-throughput sequencing and other genomic analyses and the UCSF Nikon Imaging Center for microscopy. We thank Byron Hann and the Preclinical Therapeutics core as well as the Laboratory Animal Resource Center (LARC) at UCSF. We are also grateful for the genomic data contributed by the TCGA Research Network, including donors and researchers. We acknowledge support from our colleagues at the Helen Diller Family Comprehensive Cancer Center. This work was supported by grants from the NIH (K99CA194077 and R01GM123977). SZ was supported through the HHMI Medical Research Fellowship.

## ACCESSION NUMBERS

The data for high-throughput sequencing datasets are deposited at GEO under the accession number GSEXXXX.

